# Identification of Begomovirus Genomic Components and Stress-specific Metabolic Markers in Mungbean Leaf Apoplast

**DOI:** 10.1101/2023.09.20.558570

**Authors:** Kiran Vilas Dhobale, Lingaraj Sahoo

## Abstract

Plant viruses exert control over the host metabolism to enhance their survival, but the specific sites where altered metabolites take effect remain enigmatic. This study focuses on the apoplastic region of symptomatic mungbean leaves infected with mungbean yellow mosaic India virus (MYMIV) to gain new insights into metabolite changes during infection. Leveraging NMR-based metabolome fingerprinting, we identified several stress-associated metabolites. Notably, proline and 2-Hydroxybutyrate were found to be up-regulated in the apoplast region, while down-regulated in the cytosol. Additionally, aspartate was found to be upregulated in the cytosolic region but absent in the apoplast. Importantly, our molecular analysis of the apoplast suggests the presence of MYMIV genomic components. Moreover, we characterized two distinct populations of extracellular vesicles (EVs) using ultracentrifugation, transmission electron microscopy, dynamic light scattering, and fluorometric assays. The data revealed alterations in the size and count of EVs, suggesting their potential role in facilitating the transport of viral components for long-distance cell-to-cell movement during infection. These findings provide valuable insights into the apoplast’s role and the significance of EVs in MYMIV infection, contributing to a better understanding of virus-host interactions and potentially informing new virus defense strategies.

## INTRODUCTION

The apoplast is a dynamic and essential extracellular space surrounding plant cells, playing a pivotal role in various physiological processes (Sattelmacher, 2001). It encompasses the region outside the plasma membrane, including the cell wall, xylem vessels, and intercellular spaces filled with apoplastic fluid (AF) (Sattelmacher, 2009). This complex fluid contains a diverse array of biomolecules, such as water, sugars, amino acids, secondary metabolites, DNA, RNA, cell wall-modifying enzymes, growth regulators, and stress-related proteins, making it a crucial conduit for various essential processes like water transport, photosynthesis, and nutrient distribution (Sakurai, 1998; Boudart et al., 2005; Blomster et al., 2011; Baker et al., 2012; Zand Karimi et al., 2022). Concomitantly, the apoplast’s dynamic extracellular milieu offers a hospitable niche for pathogenic colonization and proliferation, thereby hosting endophytic pathogens within plant tissues (James et al., 2001; Liu et al., 2017).

Under stress conditions, plant cells actively secrete numerous biomolecules into the AF, with a significant role played by extracellular vesicles (EVs), particularly exosomes (Pérez-Bermúdez et al., 2017; Pegtel and Gould, 2019). Exosomes, a subset of EVs, have emerged as key players in transport, cellular signaling, communication, and potentially cross-kingdom communication (Cui et al., 2020; Cai et al., 2021; Zhou et al., 2022). They have been implicated in transporting defense compounds, proteins, and small RNAs into the extracellular space, thereby facilitating inter-organism communication and enhancing plant defense mechanisms (Regente et al., 2009; Suharta et al., 2021; Alghuthaymi et al., 2021; Kim et al., 2022). A growing body of evidence indicates that cells infected with enveloped or nonenveloped viruses release EVs that contain viral components. While in animals, viruses have been shown to exploit exosomes for transmission, cloaking themselves to evade the host immune system, and facilitating long-distance cell-to-cell movement (Canitano et al., 2013; Ramakrishnaiah et al., 2013; Ahsan et al., 2016). Similar roles of apoplast and exosomes in plant-virus interactions have been demonstrated (Rutter and Innes, 2017; Movahed et al., 2019; Chen et al., 2021; Hu et al., 2021; Lu et al., 2022), yet their specific involvement in begomovirus infection remains an unexplored area of research.

The apoplastic landscape evolves significantly during pathogenic growth, with metabolic reactions and microbial processes influencing the solute composition (Geilfus et al., 2015; Du et al., 2016; Dora et al., 2022). This composition is known to vary across plant species and genotypes, responding to shifting environmental cues such as light, nutrition, and biotic and abiotic stresses (Hückelhoven, 2007; Agrawal et al., 2010; Farvardin et al., 2020). Investigating the apoplastic composition and its dynamic alterations, encompassing attributes like redox and osmotic potential, pH, nutrient/metabolite availability, and enzymatic activities, offers novel insights into plant responses to environmental stimuli (O’Leary et al., 2016a). However, the molecular profiling of the apoplast solution is complex due to its spatially structured and dynamic nature, wherein metabolites and ions may exhibit volatility, transience, or association with the cell wall and plasma membrane.

Metabolomics methodologies play a pivotal role in unveiling the intricate molecular interplay underpinning plant-pathogen interactions (Manchester and Anand, 2017). Although metabolite changes under biotic stress within diverse plant tissues, including roots, stems, and leaves, have been extensively explored using metabolomics techniques, analyzing apoplastic metabolites during pathogen interactions presents challenges owing to their low abundance and potential contamination during fluid extraction (Gentzel et al., 2019). Various methods exist for apoplast fluid extraction from different tissues (Dragišic Maksimovic et al., 2014), with the infiltration-centrifugation technique (Lohaus et al., 2001; Witzel et al., 2011; Nouchi et al., 2012; Leary et al., 2014), a well-established approach for leaves, offering scalability and compatibility with downstream biochemical and analytical methodologies.

There exists a wealth of research conducted on the altered metabolic state of entire symptomatic plant leaves during virus infection using metabolomics (Choi et al., 2006; Lionel et al., 2012; Mandal et al., 2012; López-Gresa et al., 2012; Palama et al., 2012; Plischke et al., 2012; Manchester and Anand, 2017; Villa-Ruano et al., 2018; Rossouw et al., 2019; Velásquez-Valle et al., 2020; J. Mishra et al., 2020; Maravi et al., 2022; More et al., 2022; Zhang et al., 2022). However, there remains a notable dearth of studies investigating the precise changes in the apoplast and cytoplasm region-specific metabolite composition during viral infection. Although some investigations have explored the region-specific metabolite composition changes within the apoplast during endophytic pathogen interactions (Führs et al., 2009; Baker et al., 2012; Floerl et al., 2012; O’Leary et al., 2016b; Green et al., 2020; Figueiredo et al., 2021), the specific alterations that occur during viral infection have, as of yet, remained largely unexplored.

Belonging to the *Geminiviridae* family, *Begomoviruses* comprise a group of plant viruses characterized by circular single-stranded DNA genomes (Fauquet et al., 2008; Zerbini et al., 2017; Fiallo-Olivé and Navas-Castillo, 2023). Whitefly transmission facilitates their spread, causing considerable damage to various economically significant crops globally (Czosnek et al., 2017a). A prominent member of this family is the *Mungbean yellow mosaic India virus* (MYMIV), which targets leguminous crops (Mishra et al., 2020; Qazi et al., 2007). Mungbean (*Vigna radiata* L.) stands as a crucial leguminous crop, revered for its nutritional value and nitrogen-fixing prowess, bolstering soil fertility (Pataczek et al., 2018). Among mungbean cultivars (cv.), the K851 variety stands susceptible to MYMIV infection, resulting in typical yellow mosaic disease (YMD) symptoms such as yellow mosaic patterns on leaves, leaf curling, stunted growth, and diminished yield (Maiti and Basak, 2011; Taylor et al., 2013; Bhanu et al., 2017; Kumar et al., 2021; Dhobale et al., 2023). Given mungbean’s importance as a staple food source, MYMIV infection poses a threat to food security (Karthikeyan, 2014; Kamaal and Akram, 2016).

The interaction between MYMIV and the susceptible mungbean cv. K851 offers an exemplary model system to delve into plant-virus interactions, unravelling the intricate mechanisms underlying viral pathogenesis. Scrutinizing the specific metabolic transformations within mungbean’s apoplastic and cytoplasmic domains during MYMIV infection holds the potential to unveil the viral strategies for host manipulation and evasion of plant defense mechanisms. Further, delving into the dynamics of extracellular vesicles, particularly exosomes, during MYMIV infection in mungbean cv. K851 may provide insights into the roles of these vesicles in virus transmission and intercellular communication.

To address the knowledge gaps, the present research aims to investigate the role of the apoplast during begomovirus infection in MYMIV-susceptible mungbean cv. K851 plants. We attempted to detect and identify apoplast- and cytosol-specific metabolite markers using NMR-based metabolomics. Additionally, molecular analysis techniques such as rolling circle amplification (RCA) and PCR were employed to check the presence of viral genomic components in the apoplast. Additionally, exosomes were characterized using transmission electron microscopy (TEM), dynamic light scattering (DLS), and quantified using DiOC_6_ dye.

## MATERIALS AND METHODS

### Plant selection and agroinoculation

Agro-infectious clones MYMIV DNA-A and DNA-B (Acc. no. OK431083 and OK431084) were used to agroinoculate mungbean cv. K851 (susceptible host) plants from our previous study (Dhobale et al., 2023). The abaxial surface of young trifoliate leaves of 3–4-week-old mungbean plants were infiltrated with the clones. Three biological replicates were maintained in an insect-free greenhouse and monitored for the appearance of YMD symptoms. After 30 days post infection (dpi), symptomatic leaf samples were collected and total genomic DNA (gDNA) was isolated using HiPurA plant Genomic DNA Purification Kit (HiMedia) (**Fig. S1**). RCA was performed using phi29 DNA Polymerase (Thermo Scientific) and the RCA products were monomerized by restriction digestion with unique cutter *Pst*I (**Fig. S1**). The RCA products were also subjected to PCR to confirm the presence of MYMIV DNA-A using DNA-A–specific primer set (Table S1).

### Recovery of apoplast wash fluid (AWF) and leaf without apoplast (LWA)

To obtain AWF, symptomatic leaves from virus-infected plants and non-symptomatic leaves from uninfected plants were collected at the same time of day after 30 dpi. The AWF was isolated according to (Rutter and Innes, 2017) with slight modifications. Leaves were kept in a 60 mL syringe and infiltrated with vesicle isolation buffer (VIB) containing 20 mM 2-(N-morpholino) ethanesulfonic acid (MES) pH 6, 2 mM CaCl_2_, 0.1 M NaCl. The samples were then blotted dry, transferred to a 30 mL syringe, and centrifuged in a 50 mL falcon tube at 1800 x g for 15 minutes at 4 °C. The flow-through was transferred to a fresh tube and centrifuged at 10,000 x g for 30 minutes at 4 °C to remove any large particles from the AWF. The remaining leaf tissue was labeled as leaf without apoplast (LWA). The AWF and LWA were immediately frozen with liquid nitrogen and stored at −80 °C for future use.

### NMR sample preparation

To extract metabolites from the AWF and LWA, 1 mL AWF were snap frozen using liquid nitrogen, lyophilized, and dried. The AWF sample was then resuspended in an equal volume of aqueous methanol solution (80% v/v) and vortexed for 20 seconds (Maravi et al., 2022a). The sample was centrifuged at 12000 x g for 30 minutes at 4 °C, and the supernatant was transferred to a fresh tube and evaporated using a SpeedVac concentrator (Eppendorf AG) at room temperature for 9 hours. The dried sample was then dissolved into 600 μl of D_2_O with TMS and immediately transferred to a 5 mm NMR tube (Sigma-Aldrich). Whereas, for LWA samples, 100 mg of LWA tissue was homogenized using liquid nitrogen in sterile mortal-pestle. The powder was suspended in a 1.5 mL aqueous methanol solution (80% v/v) and vortexed for 20 seconds. The sample was further extracted by ultrasonication for 5 minutes (Pulse ON: 20 sec; Pulse OFF: 10 sec) in ice-cold conditions. All experiments were performed in triplicate.

### NMR experiments

The NMR spectroscopy analysis performed as previously described (Maravi et al., 2022b). Briefly, the prepared metabolite samples were tested on Bruker Avance III 600 MHz spectrometer ((BrukerBiospin) operating at 600.17 MHz at 25 °C temperature (acquired data: free induction decay (FID) files; acquisition time: 3 sec.; spectral width: 16.0 ppm). The metabolite signals were analyzed using 1H spectrum, 2D COSY (homonuclear correlation spectroscopy) and 2D HSQC (1H-13C heteronuclear single quantum coherence spectrometry).

### Spectra processing and statistical analysis

The acquired 1H-NMR spectrum was phase- and baseline-corrected, normalized and referenced at TMS peak (0.00 ppm) using MestReNova software (Version 9.0.1, Mestrelab Research). Then performed region binning with a bin width of 0.02 ppm ranging from 0.5 to 10.0 ppm based on sum intensities method. The methanol (3.29–3.32 ppm) and water (4.80–4.92 ppm) regions were excluded from binned dataset prior to multivariate data analysis (MVDA). The SIMCA software (version 17.0, Umetrics) was used to analyze the 1H NMR binned dataset, which was normalized by Pareto scaling and log-transformation. The classification methods such as principal component analysis (PCA) and partial least squares discriminant analysis (PLS-DA) were used to explore class differences and highlight explanatory metabolites in virus-infected and non-infected samples. Orthogonal partial least squares discriminant analysis (OPLS-DA) was employed to obtain the maximum class separation and verify the results with R^2^X and Q^2^ values. The MVDA methods were validated with 100 permutations analysis and Tukey’s test was used to identify statistically significant differences (*P* < 0.05) between metabolites.

### Metabolite identification and pathway analysis

The assigned 1H NMR peaks were used to measure the concentrations of metabolites using the ChenomX NMR Suite (ChenomX Inc.), human metabolome database (HMDB) and consulting relevant literatures (Rosenblum and Tjeerdema, 2003; Choi et al., 2006; Emwas et al., 2019; Bueno and Lopes, 2020; Li et al., 2020; Mozumder et al., 2020; Ren et al., 2021; Farag et al., 2021; Villa-ruano et al., 2021). The list of significant metabolites identified by multivariate data analysis was then used to study the differences in metabolic pathways by utilizing MetaboAnalyst (https://www.metaboanalyst.ca/).

### AWF DNA extraction and molecular analysis

From AWF solution total apoplastic DNA was extracted using phenol/chloroform extraction method with slight modifications (Spada et al., 2020). The AWF DNA was subjected to RCA to amplify the circular DNA genomes of MYMIV, using TempliPhi™ DNA amplification kit (GE Healthcare). The resulting RCA product (i.e., concatemers of viral DNA) was subsequently used as a template for another RCA reaction to enrich viral genomes. In order to detect virus accumulation (DNA-A and DNA-B) in the AF of inoculated test plants, the enriched RCA products were monomerized by restriction digestion with unique cutter *Pst*I (**Fig. 4**a). The RCA products were also subjected to PCR analysis using MYMIV DNA-A and DNA-B specific primer sets (Table S1).

### Extracellular vesicles extraction

The AWF solution was passed through a 0.2 μm membrane and then subjected to centrifugation at 10,000 x g for 30 minutes at 4°C. Two distinct populations of EVs were isolated from the AWF using successive ultracentrifugation steps. The ultracentrifugation was performed using a rotor (F50L-24 x 1.5) and 1.5 mL ultra-microtube (Thermo Scientific) in the Thermo Scientific™ Sorvall™ WX+ ultracentrifuge series at 4°C. The two populations of EVs were obtained by centrifuging the AWF at 50,000 x g for 60 minutes at 4 °C (P50) and 100,000 x g for 60 minutes at 4 °C (P100). After centrifugation, the pellets were resuspended in VIB buffer for further analysis.

### Transmission electron microscopy (TEM)

To study the shapes and sizes of EVs, TEM was performed using a JEM-2100 transmission electron microscope (JEOL). Briefly, the AWF or EVs pellets were diluted by 100-fold in VIB buffer and deposited onto a 300-mesh carbon-coated copper grid (Ted Pella, Inc.). The grids were then negatively stained with 50 µl of 2% uranyl acetate, air-dried, and imaged at 80-100 kV. The resulting TEM images were analyzed using ImageJ software.

### Dynamic light scattering (DLS)

The hydrodynamic diameter and zeta potential of the isolated EVs were measured using the Litesizer 500 Particle Analyzer (Anton Paar). Each retained EVs pellet (P50 and P100) was resuspended in 700 µl of a 50 mM Tris-Cl pH 7.5 solution for DLS analysis. The particle size measurement was carried out in a polystyrene 10 x 10 x 45 mm cuvette, using the default settings for exosome analysis. The analysis allows for the determination of the hydrodynamic diameter of the EVs, which provides information about their size. Similarly, the zeta potential of the EVs was measured in an omega cuvette, following the default settings. The zeta potential is a measure of the surface charge of the EVs, which can affect their stability and interactions with other molecules.

### Fluorometric quantification of EVs

The isolated EVs quantified using a fluorometric dye (DiOC_6_) to assess total membrane content (Rutter et al., 2017). The EVs were resuspended in 100 µl of 100 µM DiOC_6_ (Invitrogen) diluted in 50 mM Tris-Cl pH 7.5. The resuspended fractions were then incubated at 37 °C for 10 minutes, washed with 2 mL of the same buffer, repelleted using an ultracentrifuge at the respective speed, and the resulting pellet was resuspended in 52 µl of fresh buffer. Next, 50 µl of each sample was transferred to a 96-well, black, clear, polystyrene plate (Corning). The fluorescence of the DiOC_6_ was then recorded using Tecan i-control (Infinite 200Pro), with an excitation wavelength of 485 nm, an emission wavelength of 535 nm, and a total of 25 flashes.

## RESULTS

### Untargeted NMR-based metabolomics discovered apoplast and cytoplasm region specific metabolites

The representative 1H NMR spectra of AWF of MYMIV-infected and uninfected mungbean leaf is presented in (**Fig. 1**a, b). Similarly, 1H NMR spectra of LWA of MYMIV-infected and uninfected mungbean is shown in (**Fig.1**c, d). The comprehensive comparative 1H NMR spectra indicated significant variation in metabolite composition between the AWF and LWA in MYMIV-infected samples (**Fig. S6**). Metabolites and their chemical shifts were correctly identified using 2D NMR spectra of COSY (**Fig. S3**), HMBC (**Fig. S4**), and HSQC (**Fig. S5**).

**Fig. 1.**
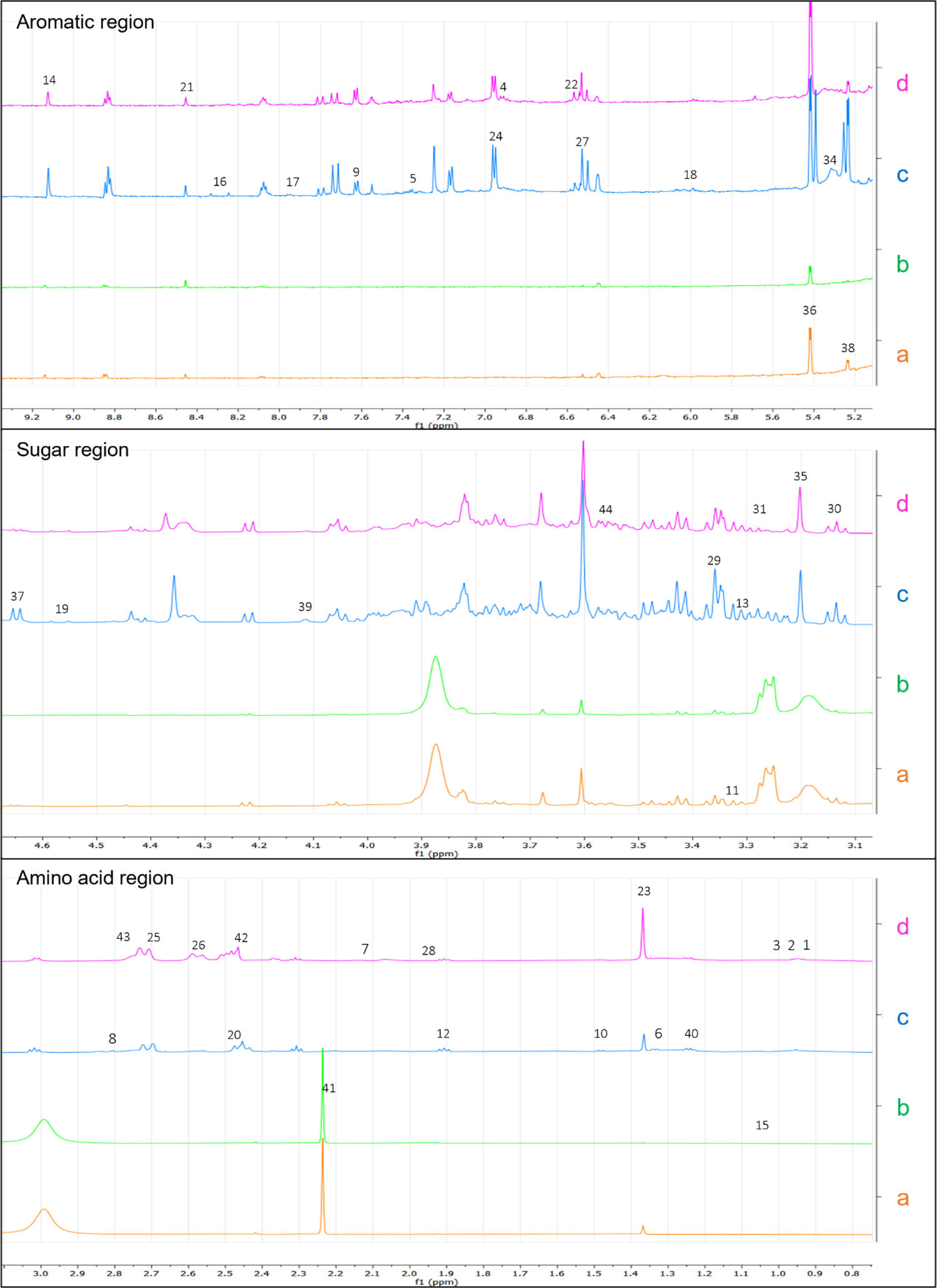
Comparative 1H NMR spectrum of MYMIV-infected and uninfected leaf tissue. a: infected AWF; b: uninfected AWF; c: infected LWA; d: uninfected LWA.

In both virus-infected and uninfected plants, approximately 21 identifiable metabolites were detected within the apoplast region. These included compounds such as α-glucose, β-glucose, sucrose, alanine, asparagine, fucose, 4-aminobutyrate, isoleucine, leucine, thionine, valine, proline, citric acid, formic acid, trigonelline, fumaric acid, betain, ethanolamine, D-ribose, pipecolic acid, and 2-hydroxybutyrate (Table 1). Moreover, in the infected apoplast region of virus-infected samples, various unidentified yet distinct metabolite signals or peaks were detected at specific chemical shift values, including 1.46 (s) ppm, 1.92 (s) ppm, 2.13 (s) ppm, 2.17 (s) ppm, 2.21 (s) ppm, 2.34 (s) ppm, 2.42 (s) ppm, 2.98 (s) ppm, 3.17 (s) ppm, 3.25 (d) ppm, 3.26 (s) ppm, 3.61 (s) ppm, and 3.87 (s) ppm.

**Table 1.** 1H NMR signals used to obtain integral regions for metabolites identification in AWF and LWA of MYMIV-infected and uninfected mungbean plants. Important features identified by fold change analysis of 1H NMR peak list using MetaboAnalyst 5.0.

| Metabolites | Chemical shifts (ppm), multiplicity, J (Hz) | AWF_<br>log2FC | LWA_<br>log2FC |
| --- | --- | --- | --- |
| Isoleucine | 1.01 (d, J=7) | NSC | NSC |
| Valine | 0.99 (d, J=7.23), 1.05 (d, J=7) | 2.737 (↑) | 1.688 (↑) |
| Leucine | 1.71 (m), 0.94 (m) | NSC | NSC |
| Tyrosine | 6.9 (d, J=8.62), 7.17 (d, J=8.65) | Absent | NSC |
| Phenylalanine | 7.36 (s), 7.43 (s) | Absent | NSC |
| Threonine | 1.33 (d) | NSC | 3.376 (↑) |
| Glutamate | 2.13 (p, J=6.86 7.29), 2.37 (dt, J=5.10, 7.92) | Absent | NSC |
| Aspartate | 2.67 (d, J=8.68), 2.81 (d, J=3.64), 2.84 (d, J=3.8) | Absent | 1.141 (↑) |
| Tryptophan | 7.63 (d, J=8.29) | Absent | NSC |
| Alanine | 1.48 (d, J=7.4) | NSC | 1.120 (↑) |
| Proline | 2.07 (m), 3.31 (m) | 1.040 (↑) | -1.05 (↓) |
| 4-Aminobutyrate | 1.91 (s), 2.31 (t, J=7.30), 3.02 (t, J=7.60) | NSC | 1.009 (↑) |
| Myo-Inositol | 3.31 (t) | NSC | 1.391 (↑) |
| Trigonelline | 9.13 (s), 8.84 (t, J=7.19, 7.19), 8.08 (t, J=7.11), 4.44 (s) | NSC | 1.624 (↑) |
| Isobutyrate | 1.08 (s) | NSC | NSC |
| Adenine | 8.25 (s) | Absent | NSC |
| Cytidine | 7.96 (d, J=8.25) | Absent | NSC |
| Adenosine | 6.07 (d) | NSC | NSC |
| β-galactose | 4.57 (d, J=19.2) | NSC | NSC |
| Succinate | 2.48 (s) | NSC | -1.054 (↓) |
| Formic acid | 8.46 (s) | NSC | NSC |
| Fumaric acid | 6.57 (s) | NSC | NSC |
| 2-Hydroxybutyrate | 1.37 (s) | 2.433 (↑) | -1.911 (↓) |
| Chlorogenate | 7.73 (d, J=15.8), 6.96 (d, J=8.1), 7.25 (s), 7.17 (d, J=8.3) | Absent | NSC |
| Citric acid | 2.74 (d, J=18.31) | NSC | NSC |
| Citrate | 2.57 (d, J=17.3) | -2.772 (↓) | -1.597 (↓) |
| Caffeoylquinic acid | 6.53 (d, J=5.65) | Absent | NSC |
| Acetic acid | 1.94 (s) | NSC | 1.388 (↑) |
| Methanol | 3.36 (s) | NSC | NSC |
| Ethanolamine | 3.13 (t, J=9.44, 9.44) | NSC | 1.816 (↑) |
| Betaine | 3.28 (s) | NSC | NSC |
| Choline | 3.2 (s) | NSC | NSC |
| Sucrose | 5.42 (d, J=3.8), 4.22 (d, J=8.8), 4.06 (t, J=8.5), 3.6 (s) | 1.130 (↑) | NSC |
| β-Glucose | 4.65 (d, J=7.95) | 2.237 (↑) | 2.933 (↑) |
| α-Glucose | 5.24 (d, J=3.7), 3.9 (dtt, J=3.43, 3.43, 7.44, 7.44, 9.59) | 2.237 (↑) | 3.222 (↑) |
| Fructose | 4.11 (d, J=3.8), 4 (t, J=11.74), 3.57 (q, J=1.99, 2.20) | NSC | 1.975 (↑) |
| Fucose | 1.24 (d, J=7.47), 1.17 (d, J=7.11) | NSC | 1.790 (↑) |
| D-Ribose | 2.2 (s) | NSC | NSC |
| Malic acid | 2.44 (d, J=11.5), 4.33 (d, J=8.1), 2.71 (d, J=17.4) | NSC | NSC |
| Shikmic acid | 2.78 (d, J=5.6), 2.75 (d, J=5.5), 6.45 (s) | Absent | NSC |
| Pipecolic acid | 3.56 (dd) | 1.479 (↑) | 1.237 (↑) |

↑: up regulated; ↓: down regulated; NSC: no significant change in metabolite; s: singlet; d: doublet; t: triplet; dd: doublet of doublets; m: multiplet; AWF: apoplast washing fluid; LWA: leaf without AWF; log2FC: log2 fold change.

Furthermore, a higher number of metabolites were identified in the cytosol (LWA) samples, which included the aforementioned 21 metabolites as well as additional compounds such as tyrosine, phenylalanine, threonine, glutamate, tryptophan, myo-inositol, caffeoylquinic acid, iso-butyrate, adenine, cytidine, shikmic acid, β-galactose, succinate, chlorogenate, acetic acid, and chlorine (Table 1). These findings indicate a substantial disparity between the metabolomic profiles of the plant leaf cytosol and their extracellular space, highlighting the distinct metabolic landscapes of these regions.

### Statistical analysis reveals significant variation in metabolite concentration and altered metabolic pathways

To study chemical differences between MYMIV-infected and uninfected mungbean plants, a statistical modeling tools like PCA, PLS and OPLS-DA were employed. Two sets of samples were analyzed: one set included the AWF of infected and uninfected plants, while the other set comprised of LWA of infected and uninfected plants.

The unsupervised method, PCA score plot of apoplast of MYMIV-infected and uninfected mungbean leaf showed good separation of the groups based on PC 1 = 37.5%, PC 2 = 22.8%, PC 3 = 16.9 % (**Fig. S2**a). Further separation between the pair confirmed via OPLS-DA model with statistical values of R^2^X = 0.755, R^2^Y = 0.996, and Q^2^ = 0.928 (**Fig. 2**a). The OPLS-DA model was validated by 200 permutations values of R^2^ = (0.0, 0.984), Q^2^ = (0.0, 0.557) (Error! Reference source not found.b). Whereas, OPLS-DA loadings S-plot was used to identify important features of the spectra that drive pair separation (**Fig. 2**c). Then we employed univariate analysis using the fold change (FC) analysis, which discriminates potentially significant peak region under study. The up-regulated metabolites in the infected apoplast were valine, 2-hydroxybutyrate, α-glucose, β-glucose, sucrose, pipecolic acid and rest of the peaks were unknown (**Fig. S7**a).

**Fig. 2.**
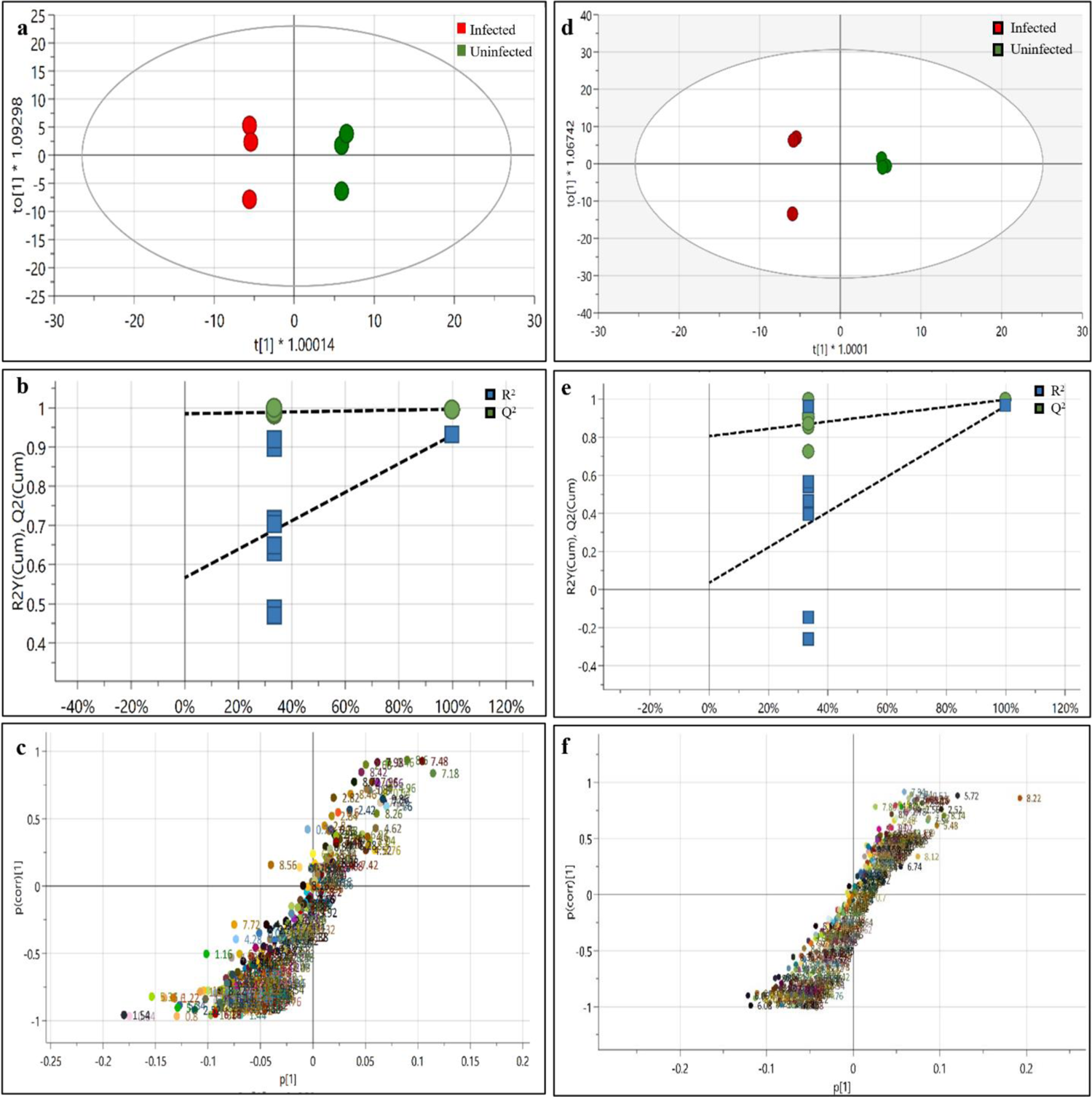
OPLS-DA score plot, statistical validation by permutation analysis and loadings S-plot for discrimination of uninfected and MYMIV-infected samples. a: Score plot of AWF samples; b: Validation by 200 permutations, R2 = (0.0, 0.985), Q2 = (0.0, 0.566) of AWF samples; c: Loadings S-plot of AWF samples; d: Score plot of LWA samples; e: Validation by 200 permutations, R2 = (0.0, 0.806), Q2 = (0.0, 0.0361) of LWA samples; f: Loadings S-plot of LWA samples.

The PCA model of LWA from MYMIV-infected and uninfected mungbean groups were separated based on PC 1 = 58.4 %, PC 2 = 22.2%, PC 3 = 17.4 % (**Fig. S2**b). The OPLS-DA model with statistical values of R^2^X = 0.921, R^2^Y = 0.997, and Q^2^ = 0.965 shown to aggregate uninfected samples and dispersed samples of infected (**Fig. 2**d). The OPLS-DA model was validated by 200 permutations values of R^2^ = (0.0, 0.836), Q^2^ = (0.0, 0.148) (**Fig. 2**e). Whereas, important features of spectra identified using OPLS-DA loadings S-plot (**Fig. 2**f). To discriminate potentially significant peak region the FC analysis was carried out. The up-regulated metabolites in the infected LWA were valine, threonine, aspartate, alanine, 4-aminobutyrate, myo-inositol, trigonelline, acetic acid, ethanolamine, α-glucose, β-glucose, sucrose, fructose, fucose, pipecolic acid and the downregulated metabolites were 2-hydroxybutyrate, citrate, proline, and succinate (**Fig. S7**b).

The concentration of selected upregulated or downregulated metabolites was measured using the ChenomX NMR Suite, and this affected list of metabolites was used for metabolite pathway analysis. The affected metabolic pathways in AWF were citrate cycle (TCA cycle), glyoxylate and dicarboxylate metabolism, arginine and proline metabolism, glycolysis / gluconeogenesis, starch and sucrose metabolism, and galactose metabolism based on the statistically significant values of (p < 0.05) and an impact factor threshold > 0 (**Fig. 3**a and Table S2). Whereas, 12 metabolic pathways were found to be affected in LWA sample i.e., inositol phosphate metabolism, phosphatidylinositol signaling system, glycine, serine and threonine metabolism, glyoxylate and dicarboxylate metabolism, glycerophospholipid metabolism, glycolysis/gluconeogenesis, sulfur metabolism, pyruvate metabolism, arginine and proline metabolism, butanoate metabolism, TCA cycle, and alanine, aspartate and glutamate metabolism (**Fig. 3**b).

**Fig. 3.**
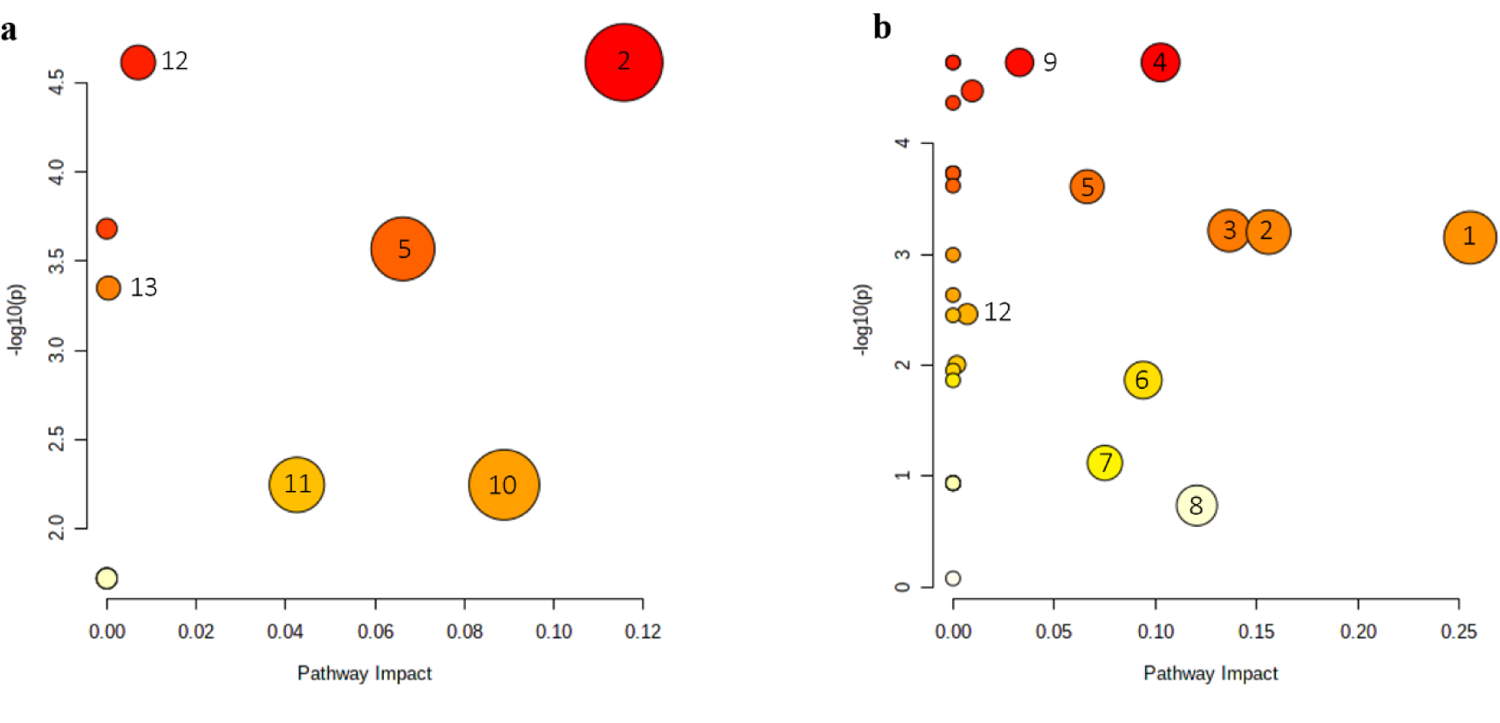
Alterations in the metabolic pathways of *Vigna radiata* cv. K851 infected with MYMIV, detected using MetaboAnalyst 5.0. Differences were considered statistically significant with P values < 0.05 and an impact factor threshold > 0. a: Altered metabolic pathways of AWF samples; b: Altered metabolic pathways of LWA samples. 1) Alanine, aspartate and glutamate metabolism, 2) Citrate cycle (TCA cycle), 3) Butanoate metabolism, 4) Inositol phosphate metabolism, 5) Arginine and proline metabolism, 6) Sulfur metabolism, 7) Pyruvate metabolism, 8) Glycine, serine and threonine metabolism, 9) Phosphatidylinositol signaling system, 10) Starch and sucrose metabolism, and 11) Galactose metabolism, 12) Glyoxylate and dicarboxylate metabolism, 13) Glycolysis / Gluconeogenesis.

### MYMIV genomic components are present in the leaf apoplast

In this study, we aimed to investigate the presence of viral genomic components in the extracellular space of MYMIV-infected plant leaves using AWF as a sample source. We extracted total DNA from the collected AWF, but the concentration was found to be very low, likely due to limited DNA circulation in AF. To overcome this challenge, we utilized the RCA technique, which generated only a few copies of viral DNA. We then used the RCA product as a template for further amplification to obtain a higher concentration of RCA product that could be used for subsequent experiments. Restriction digestion of the RCA product using the *Pst*I enzyme revealed the presence of a ∼2.7 kb band, which is indicative of the begomovirus genomic DNA molecule size, as shown in (**Fig. 4**c). To identify the presence of both genomic components of MYMIV, i.e., DNA-A and DNA-B, we performed PCR analysis using specific primer pairs.

**Fig. 4.**
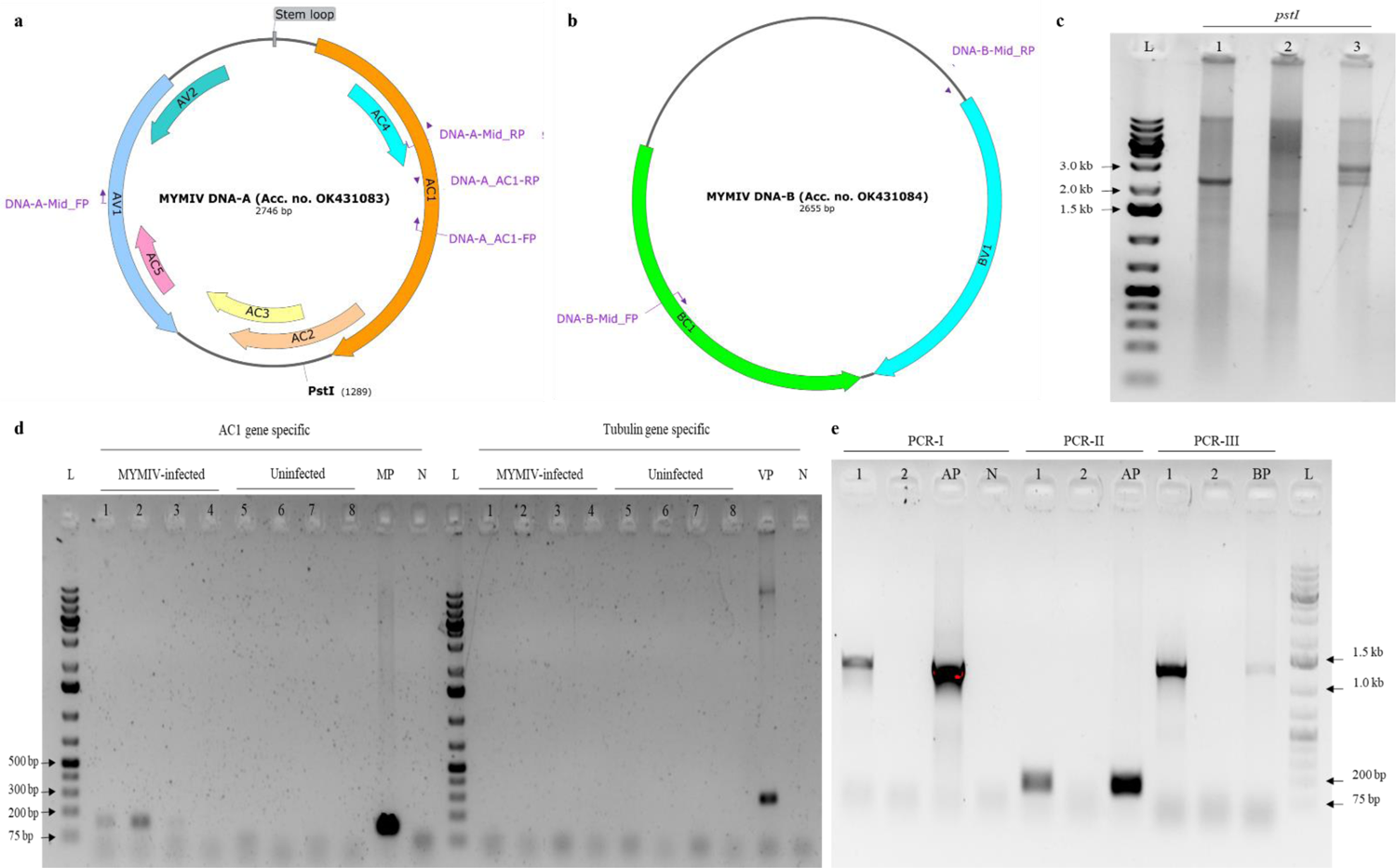
Identification of MYMIV genomic components (DNA-A and DNA-B) in the symptomatic mungbean leaf apoplast. a: Schematic representation of MYMIV DNA-A genome along with two sets of primer pairs (denoted as DNA-A_Mid and AC1_gene) for PCR detection. The unique cutter, *Pst*I, is indicated; b: Schematic representation of MYMIV DNA-B genome along with a primer pair (denoted as DNA-B_Mid) for PCR detection; c: Confirmation of the complete genome size of viral DNA (∼2.7 kb) in AWF samples of MYMIV-infected and uninfected leaf samples using unique cutter (*Pst*I) restriction digestion analysis. The following samples were used: 1: RCA product of AWF extracted from MYMIV-infected leaf-1, 2: RCA product of AWF extracted from uninfected leaf, 3: RCA product of AWF extracted from MYMIV-infected leaf-2, L: DNA ladder; d: Detection of viral DNA and host genomic DNA contamination in AWF samples of MYMIV-infected and uninfected leaf samples using PCR analysis. To specifically detect viral DNA, a primer specific to the AC1 gene of MYMIV was used. To check for any potential contamination of cytoplasmic DNA in AWF, a Vigna radiata tubulin gene-specific primer was used. PCR with the AC1 gene-specific primer produced a 137 bp fragment, while PCR with the tubulin-specific primer produced a 308 bp fragment. The following samples were used: four biological replicates (MYMIV-infected AWF sample no. 1, 2, 3, 4) and (Uninfected AWF sample no. 5, 6, 7, 8), L: 1 kb Plus DNA ladder, MP: MYMIV DNA-A_pCAMBIA3300, N: negative control, VP: total genomic DNA extracted from a healthy mungbean leaf; e: Identification of MYMIV DNA-A and DNA-B in AWF samples of MYMIV-infected and uninfected leaf samples using PCR analysis. Three sets of primer pairs (PCR-I, PCR-II, and PCR-III) were used to detect MYMIV DNA-A and DNA-B in the RCA product of AWF samples. PCR-I amplified MYMIV DNA-A mid-region (1210 bp), PCR-II amplified MYMIV DNA-A AC1 gene region (137 bp), and PCR-III amplified MYMIV DNA-B mid-region (1430 bp). The following samples were used: 1: RCA product of AWF extracted from MYMIV-infected leaf, 2: RCA product of AWF extracted from uninfected leaf, AP: MYMIV-DNA-A_pCAMBIA3300, BP: MYMIV-DNA-B_pCAMBIA3300, N: negative control, L: DNA ladder.

We detected MYMIV DNA-A using two sets of primer pairs, namely DNA-A_Mid primer (PCR-I) and AC1 primer (PCR-II), both of which generated 1.2 kb and 137 bp fragments, respectively, in infected AWF samples (**Fig. 4**e). We also identified MYMIV DNA-B using the DNA-B_Mid primer (PCR-III), which generated a 1.2 kb fragment that produced the expected size of an amplicon, as confirmed by gel electrophoresis (**Fig. 4**e). To rule out any possibility of host genomic DNA contamination in AWF DNA samples, we used a housekeeping gene, tubulin-specific primer pair, which did not show any amplification in AWF samples but showed an expected band of 308 bp in the positive control (VP) i.e., total plant leaf genomic DNA sample (**Fig. 4**d).

### TEM analysis of AWF & EVs

TEM analysis of AWF from MYMIV-infected and uninfected mungbean leaves revealed the presence of numerous circular vesicles. The size and number of vesicles were measured using ImageJ software, revealing an increase in both the total number and size of vesicles in MYMIV-infected AWF compared to uninfected AWF (**Fig. 5**). Apart from this we observed several unique but unknown structures in AWF of infected samples (**Fig. S9**).

**Fig. 5.**
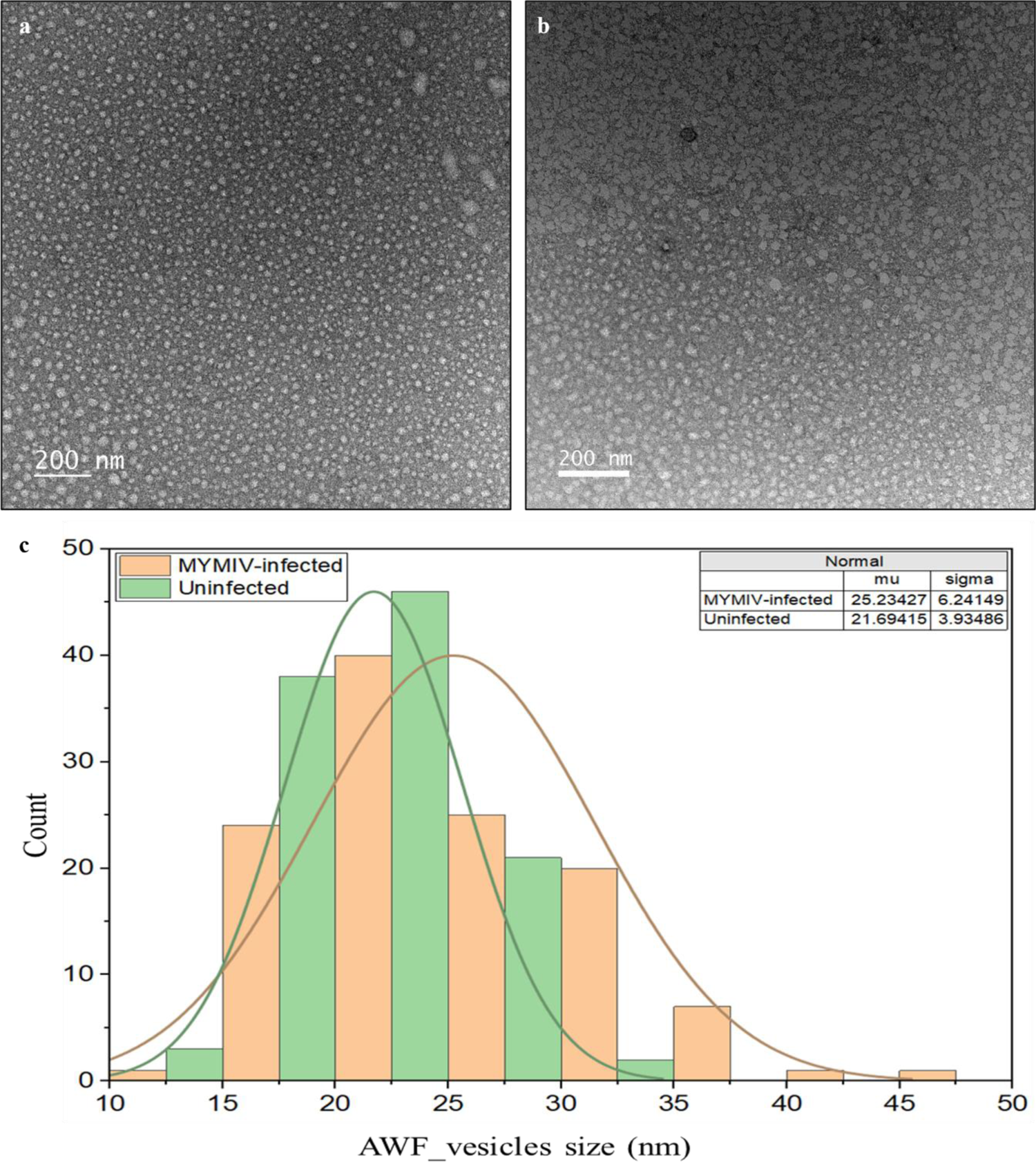
Transmission electron microscopy (TEM) image of AWF extracted from MYMIV-infected and uninfected samples. Circular shaped vesicles were visible in the apoplast fluid collected from a: MYMIV-infected and b: uninfected samples. c: vesicles size distribution graph analysis carried out using ImageJ and OriginPro software. Mu: mean size, sigma: standard deviation.

To isolate potential complete virus particles, the AWF solution underwent ultracentrifugation at 40,000 x g for 60 minutes at 4 °C, resulting in a pellet (P40). The pellet was then resuspended in VIB buffer and subjected to TEM analysis. The objective was to identify twinned quasi-icosahedral virus particles with expected dimensions of approximately 18 x 30 nanometers (Hesketh et al., 2018). However, no exact virus particles were observed. Instead, intriguing virion-like structures in a different size range of approximately 35 x 60 nanometers were detected. The TEM image in **Fig. S8** showcases these unexpected structures, prompting further investigation into their nature and significance.

### MYMIV infection cause alterations in the population of EVs

Furthermore, we employed fluorometric assays and dynamic light scattering analysis to study the concentration, hydrodynamic diameter and zeta potential of different populations of secreted EVs (P50 and P100). The P50 fraction of infected and uninfected AWF showed increased population of EVs in terms of their elevated size and concentration (**Fig. 6**a, d). Interestingly, the P100 EVs population showed a lowered concentration in infected compared to uninfected AWF (**Fig. 6**b). However, the MYMIV-infected AWF had two separate populations of EVs in the range of 70-90 and 120-140 nm, while the uninfected AWF had only single population of EVs (70-90 nm) (**Fig. 6**e). These findings suggest that MYMIV infection could have a significant impact on the population and characteristics of EVs in AWF.

**Fig. 6.**
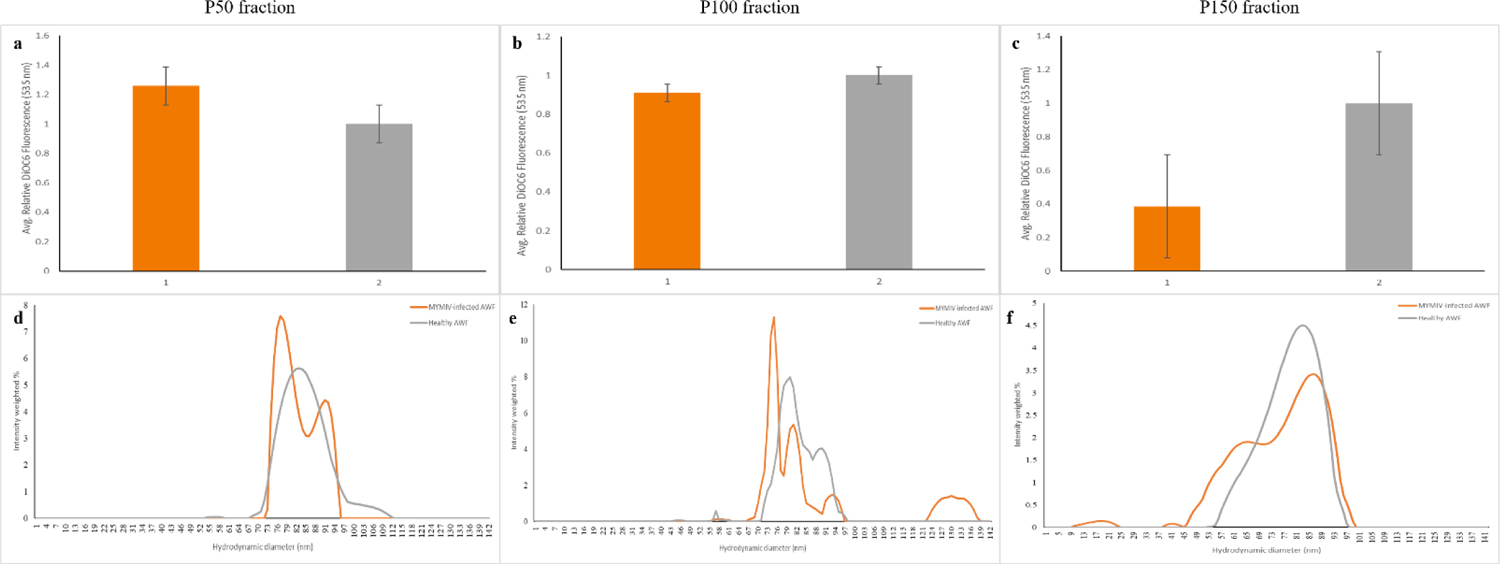
Alteration of different populations of EVs size and concentration during MYMIV infection in mungbean plants. Histograms representing the quantification of DiOC6 fluorescence in different fractions of secreted EVs. The fractions are labeled as a: P50 fraction and b: P100 fraction. The data used for this graph includes three biological replicates, each with three technical replicates. The experiment aimed to compare EV secretion between two groups: 1. EVs isolated from MYMIV-infected AWF and 2. EVs isolated from healthy AWF. Fluorescence intensity was measured, and error bars indicate the standard deviation (sd). Statistical analysis was performed using a two-tailed unpaired Student’s t-test with a significance level of P < 0.001 (n = 3). The graph represents the hydrodynamic diameter of EVs determined using DLS in different fractions. The fractions are labeled as d: P50 fraction and e: P100 fraction of secreted EVs.

## DISCUSSION

This study provides novel insights into the plant-virus interaction and sheds light on the potential role of the apoplast in begomovirus infection. Our findings suggest that the apoplast plays a crucial role in the systemic transport of viral components as evidenced by the identification of viral DNA in the AF and the elevated size and number of EVs in response to the infection. In addition, we successfully identified apoplast (AWF) and cytosol (LWA) specific metabolic biomarkers associated with MYMIV infection in mungbean plants using untargeted NMR-based metabolomics. Our results revealed significant differences in the metabolomic profiles between the apoplast region and cytosol of mungbean leaf, and MYMIV infection had a significant impact on the metabolites present in these regions.

In the present study, our findings encompass the identification and characterization of distinct EVs population within the apoplastic environment, coupled with their notable remodeling upon viral infection. These observations are in alignment with the concept that both mammalian and plant cells enhance EVs secretion during stress conditions (Hoen et al., 2016). McMullen et al., (1977) also reported proliferation of host plant EVs when infected by *Barley stripe mosaic virus*. Importantly, the present study reports the presence of both genomic components, MYMIV DNA-A and DNA-B, of the bipartite begomovirus in the apoplastic region of mungbean leaves, indicating the possibility of cell-to-cell movement via the apoplast. Plant viruses usually spread through plasmodesmata to adjacent cells by altering its size exclusion limit (Schoelz et al., 2011). This observation resonates with recent evidence suggesting that plant viruses might exploit the extracellular milieu and EVs for their dissemination, although the precise mechanisms remain to be fully deciphered (Liu et al., 2021). Insect vectors are also known to carry viral components enclosed inside exosomes for horizontal transmission to the host (Pakkianathan et al., 2015; Czosnek et al., 2017; Chen et al., 2021; Lu et al., 2022). Studies have also addressed the role of endomembrane deformation during virus replication in plants (Grangeon et al., 2012; Li et al., 2016; Agaoua et al., 2021), and the V3 protein from *Tomato yellow leaf curl virus* (TYLCV) has been identified as a cis-golgi-localized silencing suppressor required for full infection, which is also found to be cell wall and PD-localized (Gong et al., 2022). Moreover, *Turnip mosaic virus* (TuMV) uses endoplasmic-reticulum-based 6K2-tagged EVs to enclose its viral replication complex and transport it via the apoplast over long distances (Movahed et al., 2019). Kumari et al., 2016 showed the association of apoplastic ascorbate oxidase with movement protein of *Cucumber mosaic virus*.

The presence of the virus resulted in significant alterations in the apoplast metabolome. Our analysis, revealed significant differences in metabolite composition between infected and uninfected plants (**Fig. 1**). In particular, we found that AWF from infected plants contained several stress-specific metabolites alterations and a large number of unknown peaks (**Fig. 1**a, b). We identified 21 metabolites in AWF and 37 metabolites in LWA, which are common in both infected and uninfected plants (Table 1). These findings suggest that MYMIV infection significantly alters metabolite composition in mungbean leaves. Green et al., (2020) used non-targeted liquid chromatography-coupled high-resolution electrospray ionization mass spectroscopy (LC-HR-MS) to analyze the metabolome of AWF from *Epichloë*-infected and uninfected *Lolium perenne* plants. They detected endophyte-specific metabolites and a large number of unknown signals. Maravi et al., (2022) previously conducted a study on the metabolic changes in symptomatic and healthy leaves of mungbean plants infected with MYMIV. Their findings revealed overall similar metabolic changes in the leaves under viral stress, which is consistent with our results. Various metabolite markers have been identified in different studies that investigated the alteration of plant metabolome under geminivirus infection (López-gresa et al., 2012; Villa-ruano et al., 2018; Velásquez-valle et al., 2020; Donati et al., 2022). Additionally, a similar set of 1H NMR peaks for legume plant metabolome has been previously reported in several studies (Chen et al., 2019; Bueno and Lopes, 2020; Villa-ruano et al., 2021).

Furthermore, we utilized statistical modeling tools such as PCA, PLS, and OPLS-DA to investigate the chemical differences in the apoplast of MYMIV-infected and uninfected mungbean plants (**Fig. 2**). The PCA score plot showed clear separation between the groups, which was further confirmed by OPLS-DA modeling. The FC analysis revealed significant metabolites and their concentration changes (**Fig. S7**). Upregulated metabolites in the infected apoplast included valine, proline, 2-hydroxybutyrate, α-glucose, β-glucose, sucrose, pipecolic acid, and unknown metabolites, while citrate was downregulated. The higher accumulation of amino acids and their derivatives is often observed in plants infected with viruses, which is thought to be linked to stress responses triggered by the pathogen (Palama et al., 2012; Mhlongo et al., 2020). Valine and proline are two multifunctional amino acids that have been shown to confer resistance against plant pathogens (Szabados and Savouré, 2010; Anzano et al., 2022). The upregulation of proline in the apoplast and its downregulation in the cytosol suggests that proline is being transported out of the cells in response to virus infection (Table 1). Resistant plants typically accumulate higher levels of amino acids in response to pathogen infection, while susceptible plants do not (Fabro et al., 2004). The upregulation of this amino acids in the apoplast of mungbean plant leaves in response to MYMIV infection suggests that it may play a role in the plant’s defense against this virus. The 2-hydroxybutyrate is a metabolic intermediate that is involved in the production of energy. α-Glucose and β-glucose are simple sugars that are used for energy and as building blocks for other molecules. Sucrose is a disaccharide that is used for energy transport. Pipecolic acid is a non-proteinaceous product that is considered as a relevant regulator of immunity in plants and humans (Wang et al., 2018). It is a derived product of lysine catabolism. The upregulation of pipecolic acid in the apoplast of MYMIV-infected mungbean plants suggests that it may play a role in the plant’s defense against the virus. The downregulation of citrate in the apoplast of MYMIV-infected mungbean plants is also interesting. Citrate is an organic acid that is involved in the Krebs cycle, a process that generates energy in the cell (Williams and O’Neill, 2018). The downregulation of citrate suggests that the plant is diverting energy away from the Krebs cycle and towards other processes, such as the production of other metabolites.

Shikimic acid, caffeoylquinic acid, and chlorogenate are precursors to lignin, a polymer that provides strength and rigidity to plants. Lignin also helps protect plants from pathogens (Vanholme et al., 2010). Aromatic amino acids, such as tryptophan, phenylalanine, tyrosine, and glutamate, are involved in plant pathogen resistance. They are used to synthesize phenylpropanoids, which are precursors to flavonoids, monolignols, phenolic acids, and stilbenes (Rojas et al., 2014). However, our analysis of the cytosol and apoplast regions of infected plants suggests that these metabolites are only found in the cytosol region, with no significant change in concentration. This suggests that they are not being transported to the apoplast region. The significance of this finding is that it suggests that the metabolites involved in plant pathogen resistance are not being produced in sufficient quantities or are not being transported to the apoplast region in sufficient quantities could be the reason for susceptibility towards MYMIV of mungbean cv. K851. This could be due to a number of factors, such as the presence of a pathogen-induced barrier that prevents the transport of these metabolites, or the fact that these metabolites are being used for other purposes, such as the synthesis of new proteins. These observations suggest that there may be a way to increase plant resistance by improving the production or transport of these metabolites into apoplast. Further research is needed to determine the exact reason why these metabolites are not being transported to the apoplast region, and to identify new ways to improve plant resistance to pathogens.

We have shown that when a plant is infected with a virus, it increases its production of aspartate in the cytosol and was absent in the apoplast. Aspartate is an amino acid that is used to synthesize many other essential amino acids (Galili, 2011). The increased production of aspartate in the cytosol may help the plant to meet the increased demand for these other amino acids in response to virus infection. In LWA samples, several metabolites were upregulated, including valine, threonine, aspartate, alanine, myo-inositol, 4-aminobutyrate (GABA), trigonelline, acetic acid, ethanolamine, α-glucose, β-glucose, sucrose, fructose, fucose, and pipecolic acid (**Fig. S7**b). Conversely, some metabolites were found to be downregulated in LWA samples, including 2-hydroxybutyrate, citrate, proline, and succinate. The up-regulation of carbohydrate pathways such as glycolysis/gluconeogenesis, glyoxylate and dicarboxylate metabolism, starch and sucrose metabolism can be explained due to increase in some of the gene expression i.e., cwINV, HXK, and PDC as a defense response (Tadege et al., 1998; Xu et al., 2021; Rojas et al., 2014). In apoplast, α-glucose, β-glucose, and sucrose concentration was up-regulated. But in infected LWA sample sucrose level was not significantly changed and had sugars i.e., fructose, fucose, α-glucose, and β-glucose. Hence, it depicts apoplast specific transport of sucrose molecules upon virus infection in susceptible mungbean variety. Overall, the up-regulation of carbohydrate metabolism pathways in response to virus infection suggests that the plant is trying to generate more energy, building blocks, and resources to fight off the infection. The transport of sucrose out of cells and into the extracellular space may be a way for the plant to deliver sucrose to other parts of the plant where it can be used for these purposes.

In conclusion, our evidence suggests that begomovirus may also move through the apoplastic route. Genomic components of the bipartite begomovirus have been found in the apoplastic region of mungbean leaves. Our analysis using 1D NMR spectroscopy has revealed the upregulation of amino acids and their derivatives, including proline, in the apoplast of infected plants. Proline has been shown to confer resistance against plant pathogens and its upregulation in the apoplast suggests that it is being transported out of the cells in response to virus infection. The downregulation of proline in the cytosol may be a way for the plant to deliver it to other parts of the plant for defense. Our analysis has also revealed that the concentration of aromatic amino acids, which are involved in plant pathogen resistance, remained unaltered in the cytosol of infected plants, but were absent in the apoplast region. This suggests that these metabolites are not being transported to this area, potentially contributing to the increased susceptibility of infected plants to virus infection. These findings highlight the importance of the apoplast in both virus spread and plant defense mechanisms, and further research is needed to better understand the specific mechanisms involved.

**Supplementary information (SI)**

The online version contains supplementary material available

## Acknowledgements

We would like to express our sincere gratitude to Prof. Gerhard Prinsloo from the Department of Agriculture and Animal Health at the University of South Africa (UNISA) in Pretoria, South Africa, for his valuable guidance and his insightful NMR-based metabolomics seminars. We gratefully acknowledge the Central Instruments Facility (CIF) of the Indian Institute of Technology Guwahati for providing access to their NMR facility.

## Author contributions

LS and KVD designed the research; KVD carried out the literature review and execution of all experiments; KVD analyzed the data; KVD prepared the complete manuscript; LS corrected the manuscript. All authors read and approved the final manuscript.

## Availability of data and materials

Relevant data are included in this paper and its associated Supplementary Information (SI).

## DECLARATIONS

**Conflict of interest:** On behalf of all authors, the corresponding author states that there is no conflict of interest.

**Ethics approval and consent to participate:** Not applicable

**Consent to participate:** Not applicable

**Consent for publication:** Not applicable

