## Supplementary material for "Identification of Begomovirus Genomic Components and Stress-specific Metabolic Markers in Mungbean Leaf Apoplast": Sup. Info.

### <u>SUPPLEMENTARY INFORMATION</u>

#### FIGURE LEGENDS

**Fig. S1.** Representative images of mungbean leaves and molecular analysis of MYMIV infected and uninfected leaf samples: **A.** Healthy mungbean trifoliate, **B.** Symptomatic MYMIV-infected leaf, **C.** PCR analysis of infected (a), positive control (b), and uninfected (c). Restriction digestion (unique cutter, PstI) of RCA product of infected (d) and positive control (e) along with DNA ladder (m).

**Fig. S2** PCA scores plot between uninfected and MYMIV-infected: a) PCA score plot of apoplast (AWF) samples; b) PCA score plot of leaf without apoplast (LWA) samples. The explained variances are shown in brackets.

**Fig. S3** 1H-1H COSY spectrum of a representative aqueous extract of leaf without apoplast (LWA) from MYMIV-infected *Vigna* radiata cv. K851.

**Fig. S4** 1H-13C HMBC spectrum of a representative aqueous extract of leaf without apoplast (LWA) from MYMIV-infected *Vigna* radiata cv. K851.

**Fig. S5** 1H-13C HSQC spectrum of a representative aqueous extract of leaf without apoplast (LWA) from MYMIV-infected *Vigna* radiata cv. K851.

**Fig. S6** 1H NMR spectrum of a representative aqueous extract of AWF (a) and LWA (b) from MYMIV-infected mungbean leaf.

**Fig. S7** Important metabolite signal identified by volcano plot which is described by both fold change threshold (x) 2 and t-tests threshold (y) 0.1. The red circles represent features above the threshold. The further its position away from the (0,0), the more significant the feature is: **a)** AWF sample group of uninfected and MYMIV-infected; **b)** LWA sample group of uninfected and MYMIV-infected.

**Fig. S8** TEM analysis of ultracentrifuged AWF solution (P40).

**Fig. S9** Transmission electron microscopy (TEM) analysis of AWF extracted from MYMIV-infected (a, b, c, and d) and uninfected (e and f) samples revealed presence of unknown structures.

**Table S1** List of primers used in current study.

**Table S2** Metabolite pathways majorly affected by MYMIV infection in AWF and LWA regions of mungbean leaf.

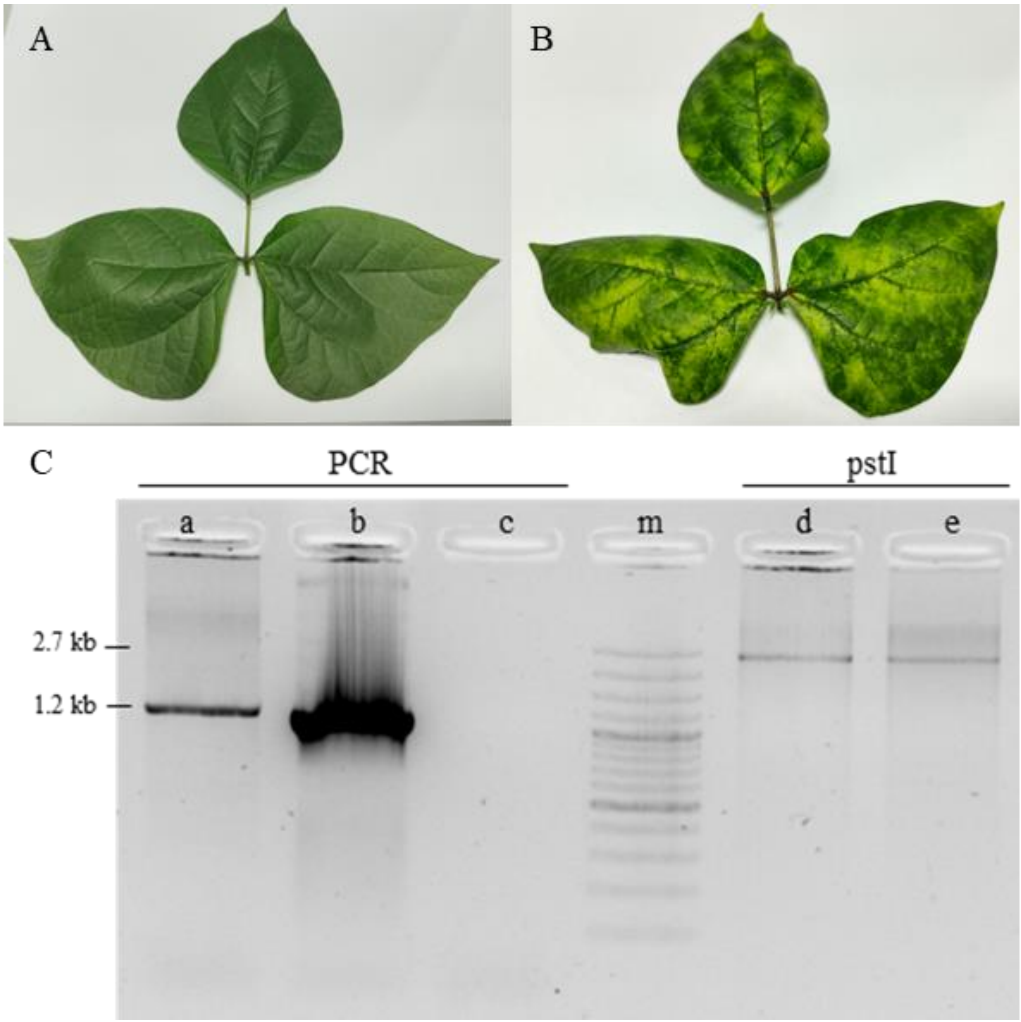


**Fig. S1** Representative images of mungbean leaves and molecular analysis of MYMIV infected and uninfected leaf samples: **A**. Healthy mungbean trifoliate, **B**. Symptomatic MYMIV-infected leaf, **C.** PCR analysis of infected (a), positive control (b), and uninfected (c). Restriction digestion (unique cutter, PstI) of RCA product of infected (d) and positive control (e) along with DNA ladder (m).

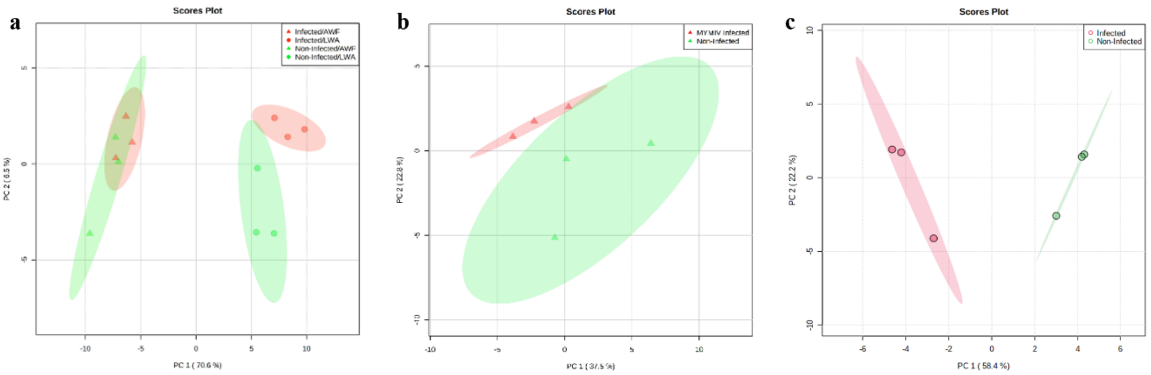


**Fig. S2** PCA scores plot between MYMIV-infected and uninfected: **a)** PCA score plot of AWF and LWA samples combined; **b)** PCA score plot of leaf without apoplast AWF samples only, **c**) PCA score plot of leaf without apoplast LWA samples only.

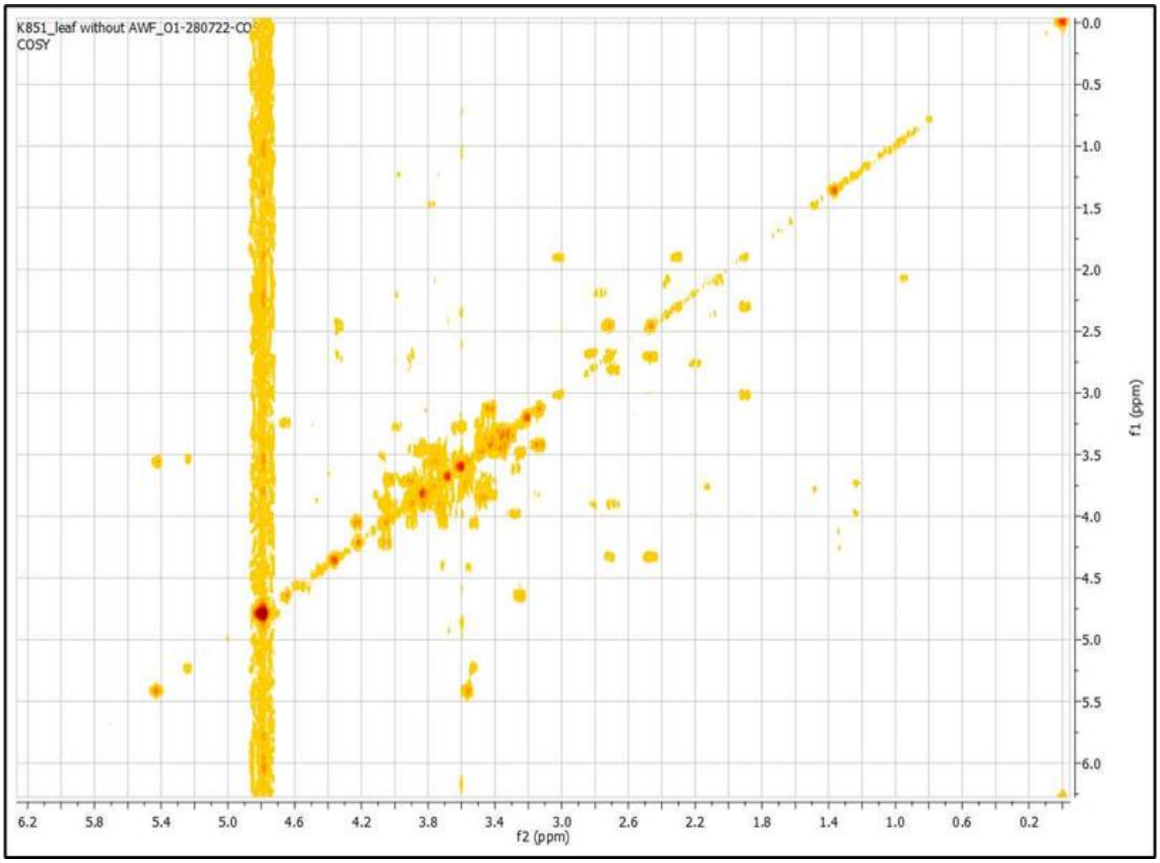


**Fig. S3** 1H-1H COSY spectrum of a representative aqueous extract of leaf without apoplast (LWA) from MYMIV-infected *Vigna* radiata cv. K851.

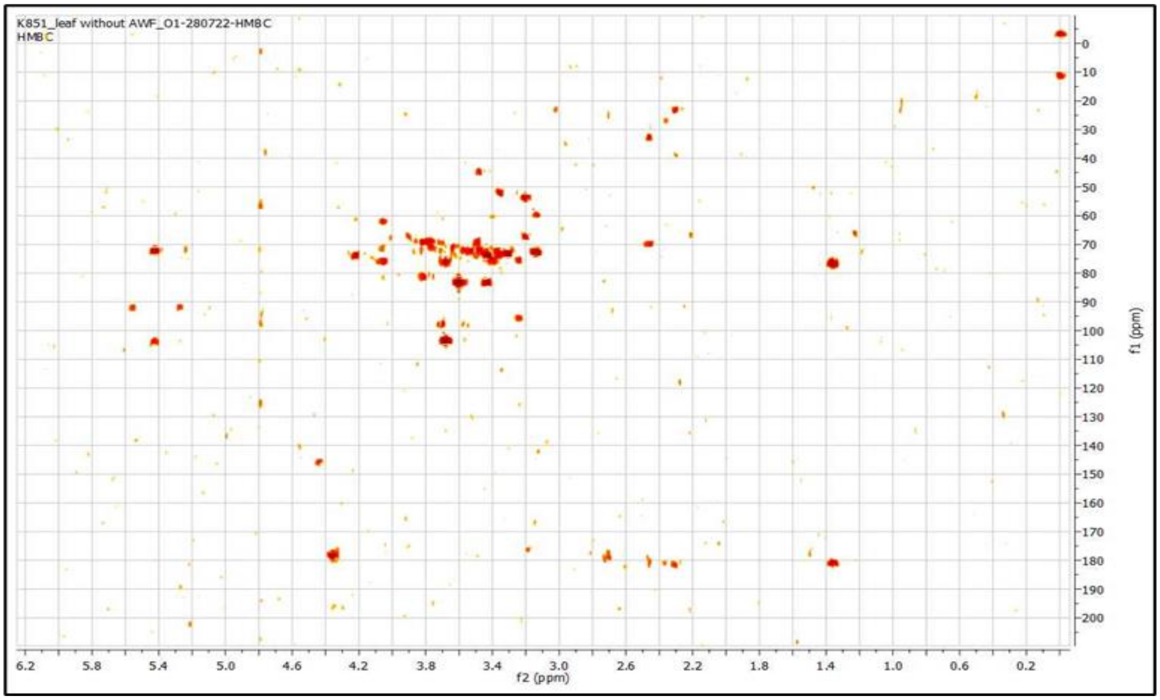


**Fig. S4** 1H-13C HMBC spectrum of a representative aqueous extract of leaf without apoplast (LWA) from MYMIV-infected *Vigna* radiata cv. K851.

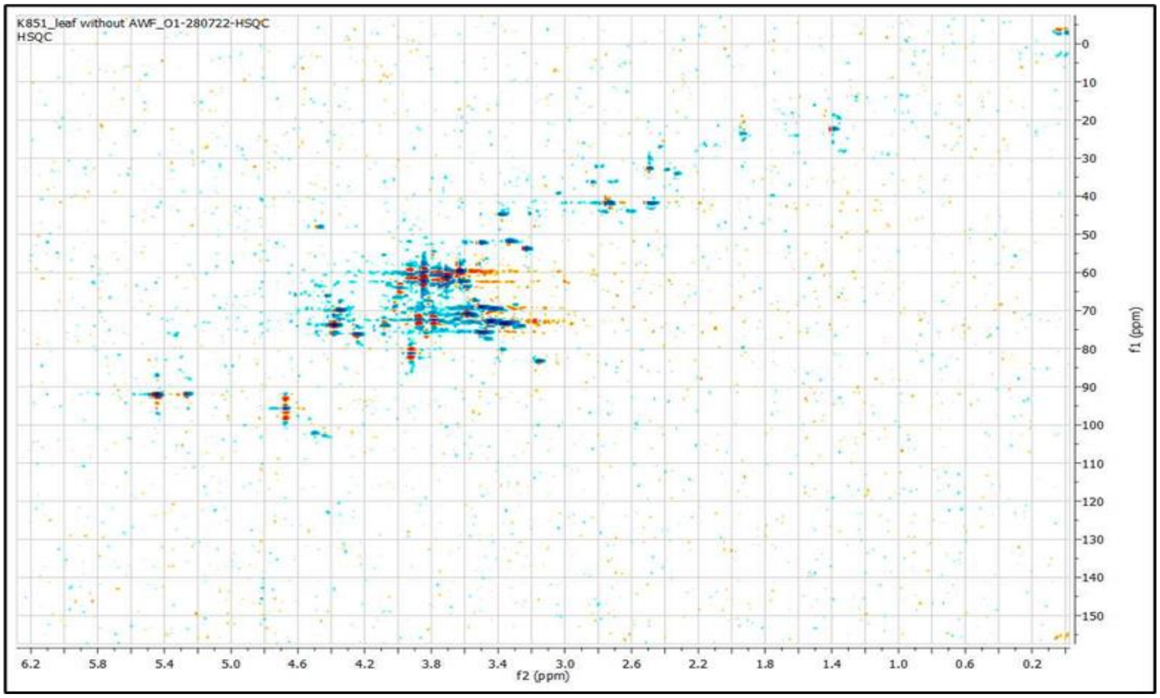


**Fig. S5** 1H-13C HSQC spectrum of a representative aqueous extract of leaf without apoplast (LWA) from MYMIV-infected *Vigna* radiata cv. K851.

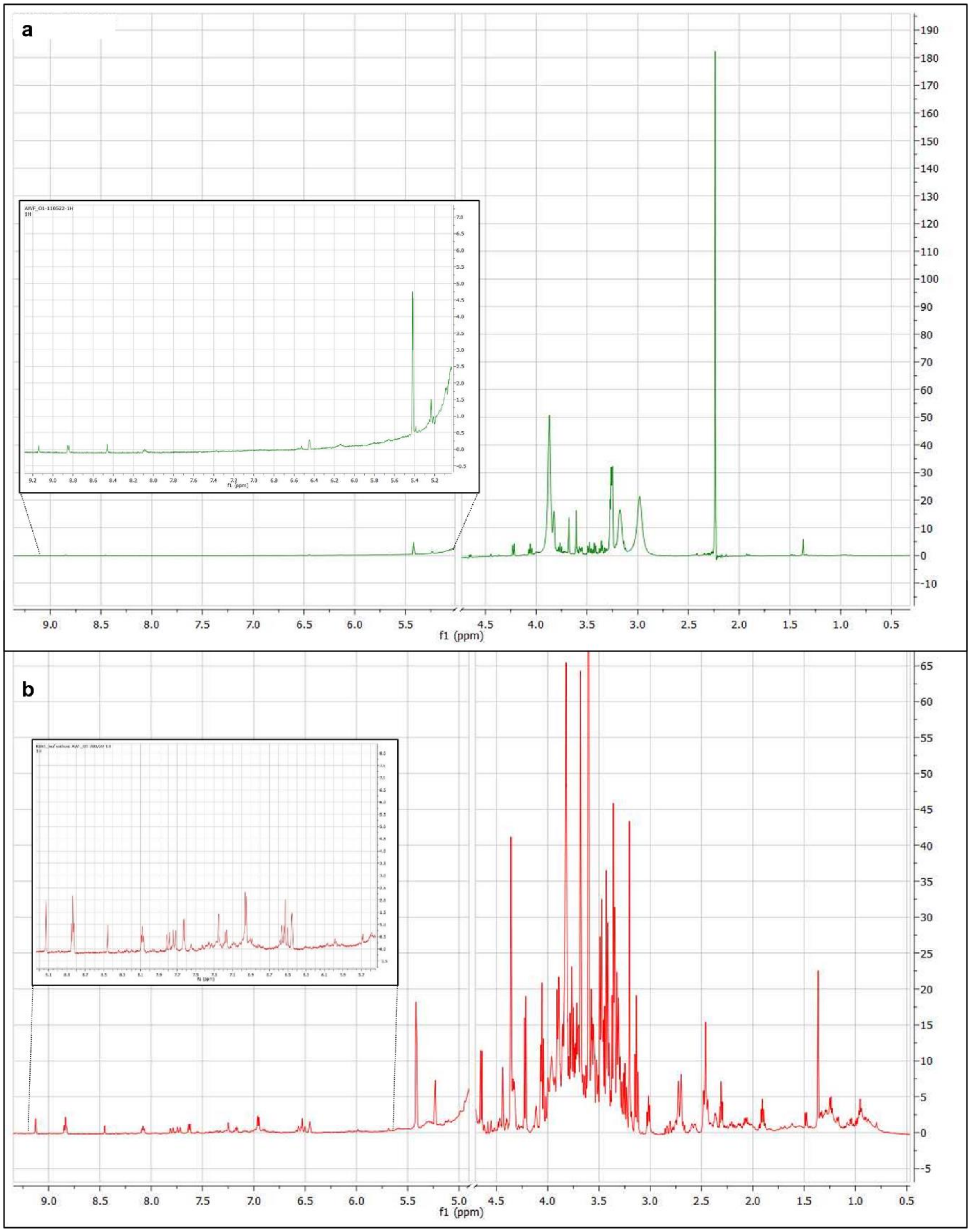


**Fig. S6** 1H NMR spectrum of a representative aqueous extract of AWF (**a**) and LWA (**b**) from MYMIV-infected mungbean leaf.

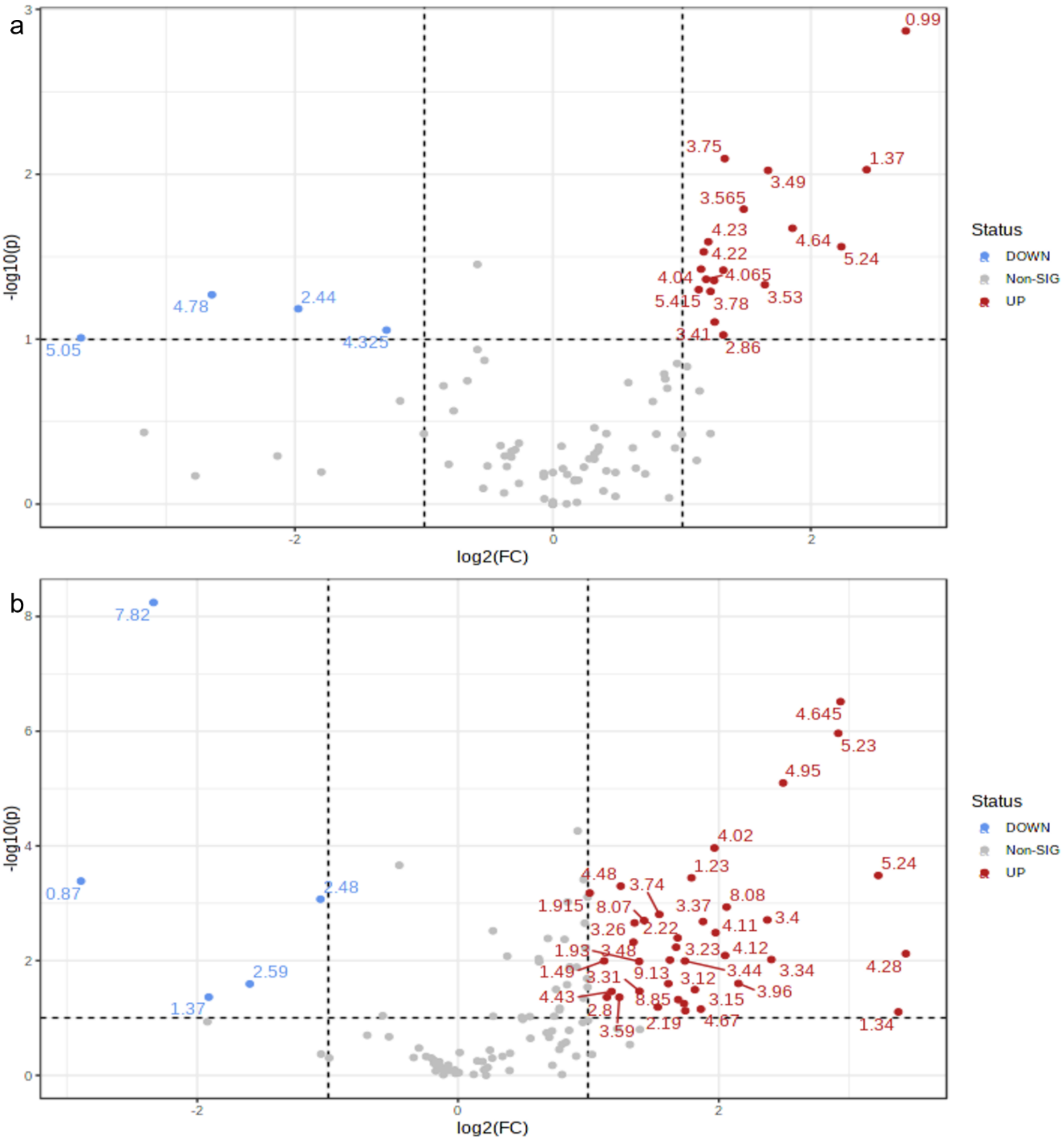


**Fig. S7** Important metabolite signal identified by volcano plot which is described by both fold change threshold (x) 2 and t-test threshold (y) 0.1. The red circles represent features above the threshold. The further its position away from the (0,0), the more significant the feature is: **a**) AWF sample group of uninfected and MYMIV-infected; **b**) LWA sample group of uninfected and MYMIV-infected.

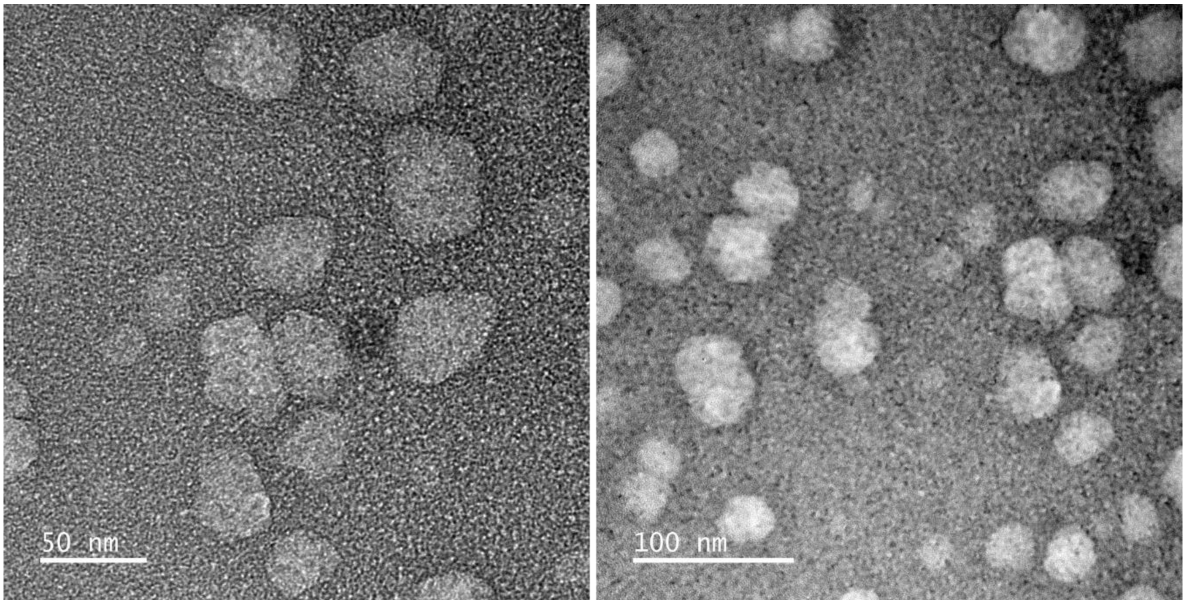


**Fig. S8** TEM analysis of ultracentrifuged AWF solution (P40).

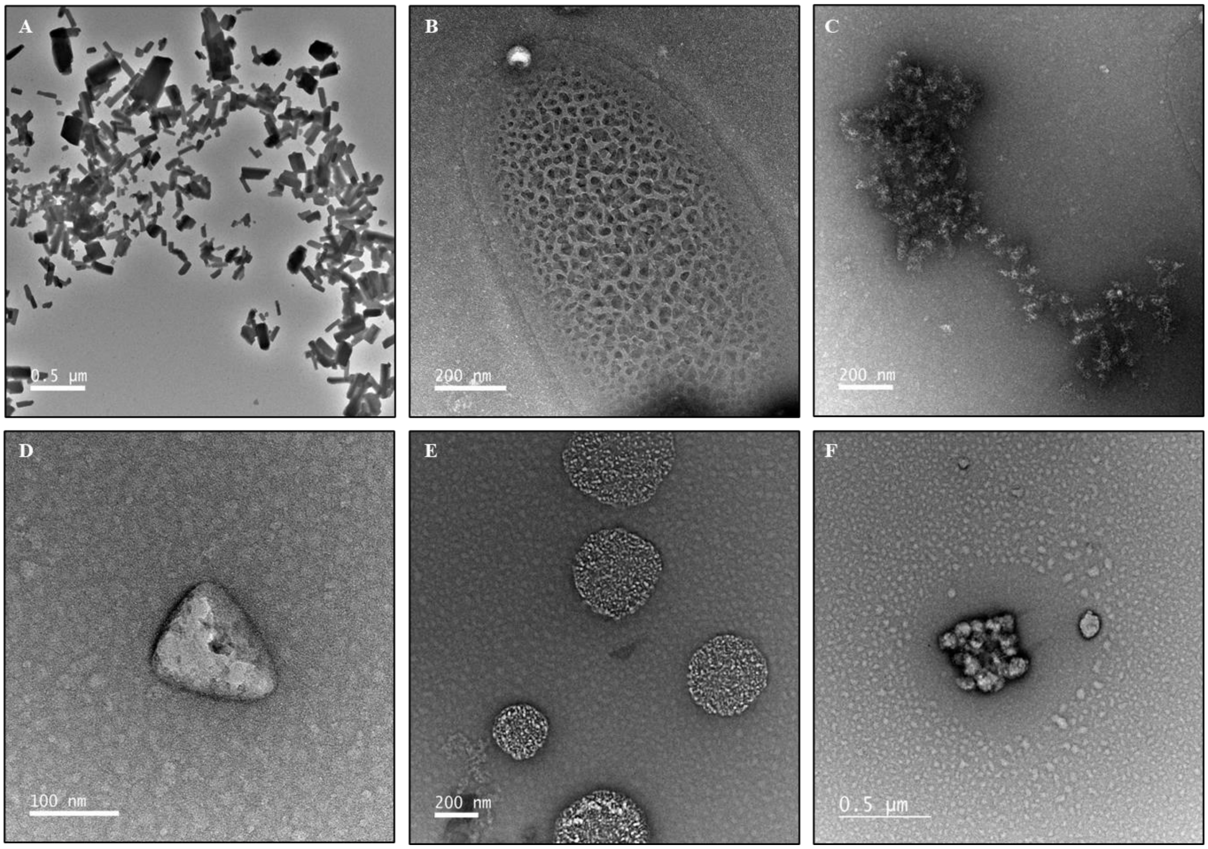


**Fig. S9** Transmission electron microscopy (TEM) analysis of AWF extracted from MYMIV-infected (**a**, **b**, **c**, and **d**) and uninfected (**e** and **f**) samples revealed presence of unknown structures.

**Table S1** List of primers used in current study.

| | Primer label | Primer pair | Sequence (5'-3') | Length | Amplicon |
|---|---|---|---|---|---|
| 1 | tubulin primer | *V*r. tubulin_FP | AACTTATCGATTCCGTCTTGGATG | 24 nt | 308 bp |
| | | *V*r. tubulin_RP | GAAGGGAAAACGGAAAACGTCATCA | 25 nt | 308 bp |
| 2 | DNA-B_Mid primer | MYMIV_DNA-B-Mid_FP | GGACTCCAATGTGATCGACGG | 22 nt | 1430 bp |
| | | MYMIV_DNA-B-Mid_RP | GCTATACGCACTATGTCGTTCG | 21 nt | 1430 bp |
| 3 | DNA-A_Mid primer | MYMIV_DNA-A-Mid_FP | CGTCCATCCATACCTTACCCG | 21 nt | 1210 bp |
| | | MYMIV_DNA-A-Mid_RP | GTATGCGTCGTTGGCAGATTG | 21 nt | 1210 bp |
| 4 | AC1 primer | MYMIV-DNA-A_AC1-FP | CTAATAGGTCTATCTGGCCGCG | 22 nt | 137 bp |
| | | MYMIV-DNA-A_AC1-RP | CGGATATTCACAGAGCCTGTCC | 22 nt | 137 bp |

**Table S2** Metabolite pathways majorly affected by MYMIV infection in AWF and LWA regions of mungbean leaf

| | **AWF Metabolic Pathway** | **Total Cmpd** | **Hits** | **Raw p** | **-log10 (*P*)** | **Holm adjust** | **FDR** | **Impact** |
|---|---|---|---|---|---|---|---|---|
| 1 | Citrate cycle (TCA cycle) | 20 | 1 | 2.43E-05 | 4.6144 | 0.00026732 | 0.00013366 | 0.11571 |
| 2 | Glyoxylate and dicarboxylate metabolism | 29 | 1 | 2.43E-05 | 4.6144 | 0.00026732 | 0.00013366 | 0.00702 |
| 3 | Arginine and proline metabolism | 34 | 1 | 0.0002703 | 3.5682 | 0.0021624 | 0.00074332 | 0.06623 |
| 4 | Glycolysis / Gluconeogenesis | 26 | 1 | 0.00044789 | 3.3488 | 0.0031352 | 0.00098536 | 0.00038 |
| 5 | Starch and sucrose metabolism | 22 | 1 | 0.0056876 | 2.2451 | 0.034126 | 0.0089377 | 0.0889 |
| 6 | Galactose metabolism | 27 | 1 | 0.0056876 | 2.2451 | 0.034126 | 0.0089377 | 0.04252 |
| | **LWA Metabolic Pathway** | | | | | | | |
| 1 | Inositol phosphate metabolism | 28 | 1 | 1.87E-05 | 4.7271 | 0.0005999 | 0.00014997 | 0.10251 |
| 2 | Phosphatidylinositol signaling system | 26 | 1 | 1.87E-05 | 4.7271 | 0.0005999 | 0.00014997 | 0.03285 |
| 3 | Glycine, serine and threonine metabolism | 33 | 2 | 0.18328 | 0.73689 | 1 | 0.18919 | 0.1204 |
| 4 | Glyoxylate and dicarboxylate metabolism | 29 | 3 | 0.0034586 | 2.4611 | 0.055337 | 0.006303 | 0.00702 |
| 5 | Glycerophospholipid metabolism | 37 | 1 | 3.38E-05 | 4.4706 | 0.00094738 | 0.00021654 | 0.00947 |
| 6 | Glycolysis / Gluconeogenesis | 26 | 2 | 0.0098468 | 2.0067 | 0.13786 | 0.016584 | 0.00189 |
| 7 | Sulfur metabolism | 15 | 2 | 0.013598 | 1.8665 | 0.16317 | 0.019868 | 0.09392 |
| 8 | Pyruvate metabolism | 22 | 1 | 0.075791 | 1.1204 | 0.75791 | 0.10545 | 0.075 |
| 9 | Arginine and proline metabolism | 34 | 2 | 0.00024644 | 3.6083 | 0.0055597 | 0.00071693 | 0.06623 |
| 10 | Butanoate metabolism | 17 | 2 | 0.00061265 | 3.2128 | 0.012866 | 0.0015569 | 0.13636 |
| 11 | Citrate cycle (TCA cycle) | 20 | 2 | 0.00063249 | 3.1989 | 0.012866 | 0.0015569 | 0.15581 |
| 12 | Ala, Asp and Glut metabolism | 22 | 4 | 0.00070787 | 3.15 | 0.01345 | 0.001618 | 0.2554 |
